# A comprehensive reference transcriptome resource for the Iberian ribbed newt *Pleurodeles waltl*, an emerging model for developmental and regeneration biology

**DOI:** 10.1101/423699

**Authors:** Masatoshi Matsunami, Miyuki Suzuki, Yoshikazu Haramoto, Akimasa Fukui, Takeshi Inoue, Katsushi Yamaguchi, Ikuo Uchiyama, Kazuki Mori, Kosuke Tashiro, Yuzuru Ito, Takashi Takeuchi, Ken-ichi T Suzuki, Kiyokazu Agata, Shuji Shigenobu, Toshinori Hayashi

## Abstract

Urodele amphibian newts have unique biological properties, notably including prominent regeneration ability. Iberian ribbed newt, *Pleurodeles waltl*, is a promising model newt along with the successful development of the easy breeding system and efficient transgenic and genome editing methods. However, genetic information of *P. waltl* was limited. In the present study, we conducted an intensive transcriptome analysis of *P. waltl* using RNA-sequencing to build gene models and annotate them. We generated 1.2 billion Illumina reads from a wide variety of samples across 11 different tissues and 9 time points during embryogenesis. They were assembled into 202,788 non-redundant contigs that appear to cover nearly complete (~98%) *P. waltl* protein-coding genes. Using the gene set as a reference, our gene network analysis identified regeneration-, developmental-stage-, and tissue-specific co-expressed gene modules. Ortholog analyses with other vertebrates revealed the gene repertoire evolution of amphibians which includes urodele-specific loss of *bmp4* and duplications of *wnt11b*. Our transcriptome resource will enhance future research employing this emerging model animal for regeneration research as well as other areas such as developmental biology, stem cell biology, cancer research, ethology and toxico-genomics. These data are available via our portal website, iNewt (http://www.nibb.ac.jp/imori/main/).

## Introduction

Urodele amphibian newts have an outstanding history as a model organism in experimental biology. The “Spemann organizer” was discovered using European newts, *Triturus cristatus* and *Triturus taeniatus*^1^. “Wolffian lens regeneration” was discovered using the newt^2^, and Eguchi et al., subsequently demonstrated transdifferentiation of pigment epithelial cells to lens cells by clonal cell culture using the Japanese fire belly newt, *Cynops pyrrhogaster*^3^. Amphibian newts have provided the clearest examples of natural reprogramming events, providing an opportunity to study mechanisms of cellular reprogramming^4-8^. Additionally, studies in newts have yielded a great deal of knowledge about the regeneration of various tissues and organs, including limb^9^, joint^10^, heart^11^, jaw ^12^, retina^13,14^, brain^15-17^, spinal cord^18^, intestine^19^, testis^20,21^, and lung^22^. Among vertebrates, only newts are known to be capable of regenerating all of the above-mentioned organs and body parts. Furthermore, comparative studies of regeneration ability between newts and frogs have provided new insights for future regenerative medicine, given that frogs (like mammals) lose the ability to regenerate various tissues and organs after metamorphosis^10,12,23,24^.

Newts also have been employed in research other than that on regeneration, reflecting these animals’ unique biological properties. The genome sizes of newt species are 8-10 times larger than the human genome^24-26^. Newts are tumor-resistant, despite having a long lifetime^27,28^. Newt eggs are fertilized via physiological polyspermy^29^. Male newts form new testes even after sexual maturation^30^. Moreover, the mating behavior of newts is mediated by sexual pheromones^31,32^. Finally, several groups have shown the utility of the newt for the toxicity testing of chemical compounds ^33-35^. Indeed, aquatic tetrapods like newts can serve as an important indicator of the influence of chemical compounds on the environment^36^. Therefore, the newt is a versatile model animal that can be used in various fields of research, including regeneration, stem cell biology, cancer research, developmental biology, reproductive biology, evolution, ethology, and toxico-genomics.

Although these properties make newt an attractive model animal, the newt species that have been used (e.g., the American common newt, *Notophthalmus viridescens*, and the Japanese common newt, *C. pyrrhogaster*) in classic experiments are not suitable for reverse or molecular genetics because of the difficulty of breeding these species in captivity. For example, Japanese common newts spawn seasonally, and each female spawns only a small number of eggs per cycle^37,38^. Three or more years can be required for sexual maturation in general. In addition, different newt species have been used for the studies performed in different laboratories, countries, and continents, making it difficult for members of this research community to share resources and expertise.

The Iberian ribbed newt (*Pleurodeles waltl*) is an emerging model newt^24,39^. In contrast to conventional newt species, *P. waltl* has a vigorous reproductive capacity and is easy to maintain in the laboratory. Using the *P. waltl* newts, we have established a model experimental system that is available for molecular genetics^39^. Notably, efficient genome editing was recently demonstrated in *P. waltl* ^40,41^. Despite the excellent utilities, the genetic information of Waltl was limited. Recently, Elewa et al. described a draft sequence of the 20-Gb giant genome and the transcriptome of *P. waltl*^24^. These data provided pioneering references for the newt research field. However, there remained a popular demand to improve the gene catalogues of this species that can be shared in the community.

In the present study, we sought to create a reference set of comprehensive gene models for *P. waltl*. To that end, we prepared 29 libraries of mRNA from various tissues and embryonic stages of *P. waltl* and subjected the libraries to RNA-seq. We combined the resulting > 1.2 billion reads and assembled these reads. This assembly yielded 1,395,387 contigs and permitted the annotation of 202,788 genes. Using these data, we built a new gene model of *P. waltl*; the resulting model expected to cover 98% of *P. waltl* protein-coding genes. Moreover, we demonstrated that the expression patterns of regeneration-specific, developmental-stage-specific, and tissue-specific genes could be analyzed using our gene model and transcriptome data sets. Finally, we have established a portal website that provides the research community with access to our data sets.

## Materials and methods

### Animals

The *P. waltl* used in this study were raised in a closed colony that originated at Tottori University. The animals were maintained as described previously^39^. The developmental stages (St) were defined according to criteria described Shi and Boucaut^42^. To isolate organs or perform surgical operations for limb and heart regeneration, embryos and adults were anesthetized/euthanized by immersion in 0.01-0.2% MS-222 (tricaine; Sigma-Aldrich, MO, USA). All procedures were carried out in accordance with Institutional Animal Care and Use Committee of the respective institute and the national guidelines of the Ministry of Education, Culture, Sports, Science & Technology of Japan.

### RNA preparation, library construction, and RNA sequencing

Sequence data collection was performed in 5 laboratories. Methods for total RNA extraction, library preparation, and sequencing of the resulting libraries are summarized in Table 1 and Supplementary Table 1.

**Table 1.** Summary of the sample preparation and sequence profiles.

| label* | tissue/organ | age/ dev.<br>stage** | description | platform<br>(Hiseq) | read<br>length# | total reads |
| --- | --- | --- | --- | --- | --- | --- |
| HtN | heart | adult | ventricle, normal | 2000 | 101 bp | 55,882,914 |
| HtR | heart | adult | ventricle, regenerating | 2000 | 101 bp | 50,516,110 |
| Lb0 | limb | adult | limb, normal | 2000 | 101 bp | 55,246,408 |
| Lb3 | limb | adult | limb, regenerating day3 | 2000 | 101 bp | 47,286,228 |
| Lb19 | limb | adult | limb, regenerating day19 | 2000 | 101 bp | 64,373,580 |
| Tl | tail | adult | tail | 2500 | 125 bp | 27,630,846 |
| Br | brain | adult | brain | 2500 | 125 bp | 27,626,778 |
| Kn | kidney | adult | kidney | 2500 | 125 bp | 27,441,344 |
| Lv | liver | adult | liver | 2500 | 125 bp | 30,427,600 |
| Pc | pancreas | adult | pancreas | 2500 | 125 bp | 24,594,822 |
| It | intestine | adult | intestine | 2500 | 125 bp | 28,214,656 |
| Cn | connective<br>tissue | 3 months<br>juvenile | connective tissue adjacent to testis | 2000 | 106 bp | 34,403,376 |
| Tg | testicular<br>grand | adult | testicular grand, matured | 2000 | 106 bp | 33,182,908 |
| Tt3 | testis | 3 months<br>juvenile | testis, not matured | 2000 | 106 bp | 36,052,648 |
| TtA | testis | adult | testis, matured | 2000 | 106 bp | 35,305,096 |
| Ov3 | ovary | 3 months<br>juvenile | ovary, not matured | 2000 | 106 bp | 37,990,436 |
| Ov7 | ovary | 7 months<br>juvenile | ovary, not matured | 2000 | 106 bp | 39,617,046 |
| Eg-1 | whole | unfertilized egg | biological replicate 1 | 2000 | 106 bp | 35,758,440 |
| Eg-2 | whole | unfertilized egg | biological replicate 2 | 2000 | 106 bp | 38,172,320 |
| St7 | whole embryo | stage 7-7.5 | early gastrula | 2500 | 125 bp | 29,361,580 |
| St8b | whole embryo | stage 8b | slightly advanced early gastrula | 1500 | 100 bp | 65,423,479 |
| St11 | whole embryo | stage 11 | middle-late gastrula | 2500 | 125 bp | 31,400,058 |
| St12 | whole embryo | stage 12 | late gastrula; | 1500 | 100 bp | 55,684,175 |
| St15 | whole embryo | stage 15 | neural plate stage | 1500 | 100 bp | 48,604,306 |
| St18 | whole embryo | stage 18 | late neural fold stage | 1500 | 100 bp | 55,446,723 |
| St25-1 | whole embryo | stage 25 | elongated tail bud stage | 1500 | 100 bp | 49,345,987 |
| St25-2 | whole embryo | stage 25 | tail bud stages (st25-28) were mixed. | 2000 | 101 bp | 51,692,566 |
| St30-1 | whole embryo | stage 30 | gill protrusion stage | 1500 | 100 bp | 49,079,133 |
| St30-2 | whole embryo | stage 30 | st30 and st31were mixed. | 2000 | 101 bp | 49,920,498 |

\*: abbreviations correspond to the labels in Table1, Fig. 3, Fig.5, Fig.7 and Supple. Fig.2 and Supple. Fig.3 #: paired end.

### Assembly and ORF prediction

All sequenced reads were employed for *de novo* assembly using the Trinity program ver. 2.4.0^43^ under default parameter settings; the trimming option was performed using the trimmomatic software^44^. Assembled contigs were processed with the TransDecoder program ver. 3.0.1^45^ to predict open reading frames (ORFs) and amino acid sequences. We used BLASTP and Pfam options for ORF predictions. To retain proteins of short length (e.g., neuropeptides), we kept ORFs of more than 50 amino acids. Redundant ORFs were filtered using the CD-HIT program^46^. The quality of the assembly was evaluated by the BUSCO program ver. 2^47^ against a core-vertebrate gene (CVG) data set^48^ and a vertebrate data set (Vertebrata_odb9).

### Gene annotation and ortholog analysis

We searched homologs of predicted amino acid sequences using a BLASTP search against the NCBI non-redundant database (nr DB) [parameters: BLAST+ ver. 2.6.0; the nr DB was the latest version as of Nov 23, 2017]. Gene ontology (GO) terms for each sequence also were annotated using the BLAST2GO program (Version 4.1.9) with the NCBI nr DB^49^. To identify vertebrate orthologs of each newt amino acid sequence, orthologous groups within vertebrates were inferred using the OrthoFinder2 program (version 2.0.0)^50^. In this ortholog analysis, we used 10 vertebrate species: green anole (*Anolis carolinensis*), zebrafish (*Danio rerio*), chicken (*Gallus gallus*), human (*Homo sapiens*), coelacanth (*Latimeria chalumnae*), mouse (*Mus musculus*), Iberian ribbed newt (*P. waltl*), Chinese softshell turtle *(Pelodiscus sinensis*), African clawed frog (*Xenopus laevis*), and western clawed frog *(Xenopus tropicalis*). All of the protein sequences were downloaded from OrthoDB ver. 9.1, except that *X. laevis* sequences were obtained from Xenbase (http://www.xenbase.org/other/static/ftpDatafiles.jsp).

### Expression and network analysis

We quantified expression of each gene in each sample by mapping to the reference transcript database that was created by the *de novo* assembly (see above). The kallisto program v0.43.1 with 100 bootstrap replicates was used for mapping and counting^51^. The read count data were normalized by the trimmed mean M values (TMM) method available in the edgeR software package of R language (version 3.12.1)^52^. After TMM normalization, we estimated the Reads Per Kilobase of exon per Million mapped reads (RPKM) value of each gene. We used RPKM values for the gene network analysis. To visualize profiles of gene expressions, the multi-dimensional scaling (MDS) plot was generated in the edgeR software package.

To detect modules of co-expressed genes among our sequencing data, weighted correlation network analysis (WGCNA) was applied. This method can identify co-expressed modules (e.g., tissue-specific gene groups) in huge data sets. Normalized RPKM data were utilized for this analysis, implemented in the WGCNA library of R language (version 1.51)^53^ with specific parameter settings of power = 8, minModuleSize = 30, and maxBlockSize = 10000.

#### Identification of *bmp2/4/16*

To identify *P. waltl bmp* genes and infer phylogeny of this gene family among vertebrates, the corresponding predicted protein sequences were used to search the genome database described below. We searched the Ensembl database version 91^54^ to identify *bmp* orthologs in 8 vertebrate species, and *X. tropicalis* v9.0 gene model in the Xenbase were also searched^55^. Additionally, we searched three independent urodele amphibian-specific data sets that were described in previous reports, including those for *Nanorana parkeri^56^, Ambystoma mexicanum^51^*, and *C. pyrrhogaster*^58^. Orthologous sequences encoded by *bmp* genes were aligned using the MUSCLE algorithm with default settings in the MEGA7 software^59,60^. A phylogenetic tree was constructed using the maximum-likelihood (ML) analysis implemented in MEGA7 with the JTT model and gamma distribution. Bootstrap probabilities were computed using 1,000 replicates.

## Results and Discussion

### Collection and preparation of material

We sought to create a comprehensive transcriptome reference covering the *P. waltl* gene repertoire, with the hope the resulting database will be useful for various subsequent studies. Therefore, we collected RNA samples from a wide variety of tissues and developmental stages (Fig. 1 and Table 1). The 29 resulting libraries were derived from 11 different normal tissues (heart, limb, brain, kidney, pancreas, tail, testicular connective tissue, testis, testicular gland, and ovary) and two regenerating tissues (heart and limb) of adult newts or, from whole embryos at each of 9 time points from early to late developmental stages (unfertilized egg and stages 7-7.5, 8b, 11, 12, 15, 18, 25, and 30).

**Figure 1.**
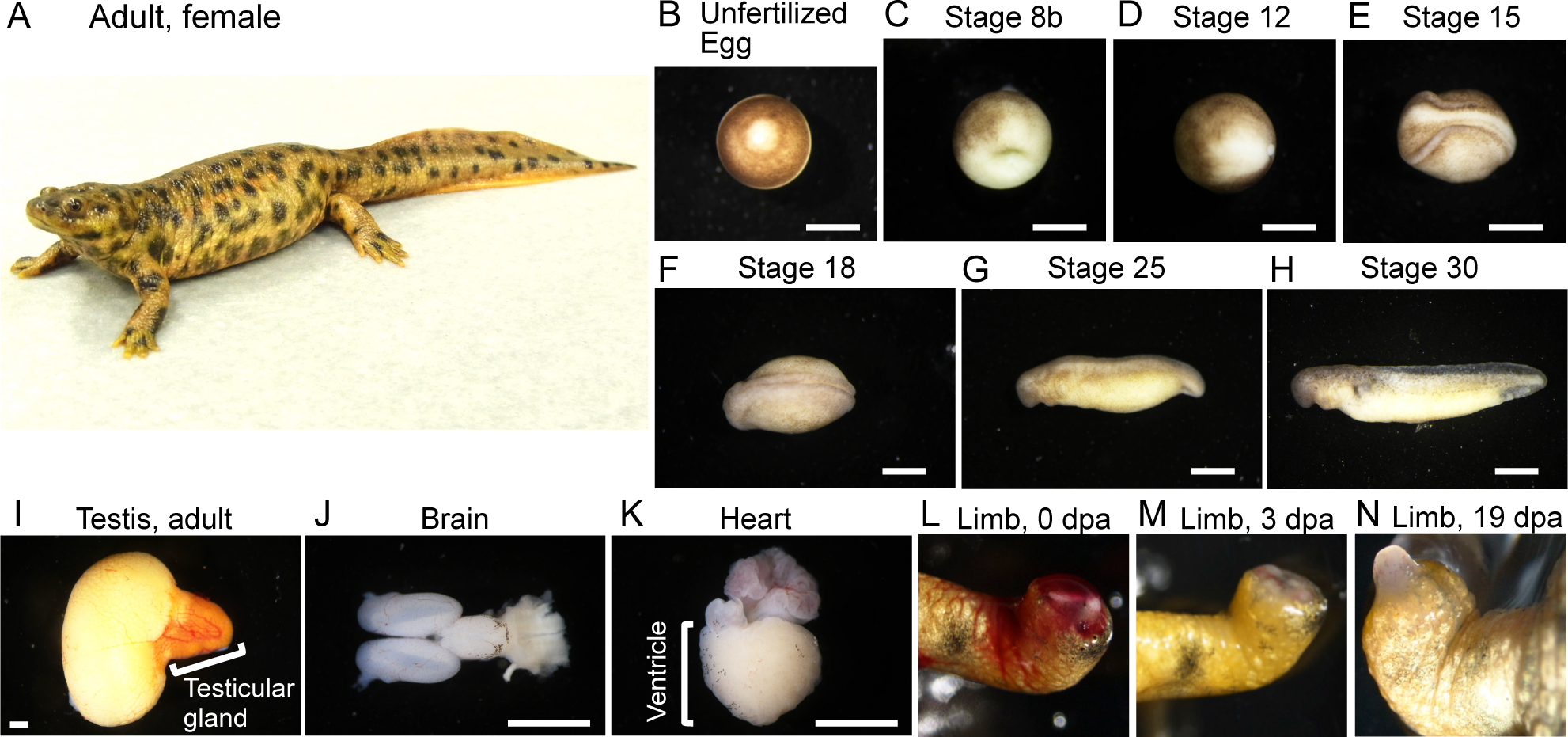
Organs and embryos used for RNA preparation. Panel (A) provides a picture of a whole adult female. (B-N) Examples of tissues and organs used for preparation of RNA. dpa: days post amputation. Scale bars: 1 mm.

### Sequencing and *de novo* assembly of transcriptome

We sequenced the 29 libraries, each of which yielded 24 to 65 million of 100- to 125-base paired-end reads, totaling more than 1.2 billion reads. To build a reference of *P. waltl* transcriptome, cleaned reads from all of these libraries were assembled together using Trinity, yielding 1,395,387 contigs with an average length and N_50_ of 700.56 bp and 1,490 bp, respectively (Table 2, Supplementary Table 2). From these contigs, we predicted 202,788 non-redundant ORFs, ranging from 147 bp to 37.1 kb with an N_50_ of 591 bp (Table 2, Supplementary Table 2). The ORF set was designated PLEWA04_ORF and used as a reference *P. waltl* coding-sequence catalogue for downstream analysis.

**Table 2.** Over view of de novo assembry and ORF prediction.

|  |  |
| --- | --- |
| **Number of samples** | 26 |
| **de novo assembly** |  |
| Total length | 977,554,621 bp |
| N50 | 1,490 bp |
| Number of contigs | 6440242 |
| **ORF prediction** |  |
| Total length | 113316939 bp |
| N50 | 591 bp |
| Number of proteins | 202788 |
| **BUSCO completeness\*** | 99.0% |

\*: including 1% of fragmented.

We evaluated the completeness of our transcriptome by comparison (via the BUSCO program) with two different datasets (CVG and Vertebrata_odb9). The CVG data consists of 233 genes that are shared as one-to-one orthologs among 29 representative vertebrate genomes and are widely used for phylogenomic studies^61^. We found that our *P. waltl* transcriptome covered all 233 CVG genes, indicating that we successfully reconstructed most of the protein-coding gene sequences in this species. In addition, our *P. waltl* transcriptome corresponded to 98% of the Vertebrata_odb9 gene set. We compared our result with earlier urodele transcriptome studies. Previous *P. waltl* and *A. mexicanum* transcriptomes covered 82% and 88% of the Vertebrata_odb9 data, respectively ^24,57^. Thus, our *P. waltl* transcriptome data significantly enhanced the gene space of urodeles, attaining a near-complete gene repertoire.

### Gene annotation and ortholog analysis

All translated sequences of PLEWA04_ORF were compared with the NCBI non-redundant protein database (nr DB) using BLASTP. Among the 202,788 ORFs identified in our DB, 121,837 genes (60.1%) encoded proteins exhibiting sequence similarity to proteins in the NCBI nr DB (Supplementary Data 1: https://doi.org/10.6084/m9.figshare.c.4237406.v1). The two most-frequent BLASTP top hit species corresponded to clawed frogs (*X. tropicalis* and *X. laevis*), followed by coelacanth (*L. chalumnae*) and turtles (*C. picta*, *P. sinensis*, and *C. mydas*) (Table 3). We used InterProScan to query the predicted coding regions for known functional domains. We identified 90,471 Pfam motifs (Supplementary Data 2: https://doi.org/10.6084/m9.figshare.c.4237406.v1) in the products of 55,075 *P. waltl* gene models. In addition, 814,803 GO terms were assigned to 86,516 genes (42.7%) (Supplementary Data 3: https://doi.org/10.6084/m9.figshare.c.4237406.v1).

**Table 3.** Species of top BLAST hits in NCBI nr databases for the *P. waltl* transcriptome

| **species** | **common name** | **# top hits** |
| --- | --- | --- |
| *X. tropicalis* | western clawed frog | 12199 |
| *X. laevis* | African clawed frog | 8604 |
| *L. chalumnae* | coelacanth | 7941 |
| *C. picta* | western painted turtle | 7631 |
| *P. sinensis* | Chinese soft-shelled turtle | 5200 |
| *C. mydas* | green sea turtle | 4229 |
| *Nanorana parkeri* | Nanorana parkeri | 3347 |
| *Larimichthys crocea* | large yellow croaker | 2827 |
| N/A | N/A | 2732 |
| *Shewanella frigidimarina* | Shewanella frigidimarina | 1960 |
| *A. carolinensis* | green anole | 1937 |
| *Alligator mississippiensis* | American alligator | 1632 |
| *Cyprinus carpio* | common carp | 1584 |
| *Oncorhynchus mykiss* | rainbow trout | 1443 |
| *Cordyceps militaris* | Cordyceps militaris CM01 | 1395 |
| *Gekko japonicus* | Gekko japonicus | 1360 |
| *Plasmodium malariae* | Plasmodium malariae | 1350 |
| *Stylophora pistillata* | Stylophora pistillata | 1323 |
| *Crassostrea virginica* | eastern oyster | 1286 |
| *Strongylocentrotus purpuratus* | purple sea urchin | 1013 |
| *Austrofundulus limnaeus* | Austrofundulus limnaeus | 978 |
| *Oreochromis niloticus* | Nile tilapia | 923 |
| *Pogona vitticeps* | central bearded dragon | 857 |
| *Daphnia magna* | Daphnia magna | 850 |
| *H. sapiens* | human | 791 |
| *Arabidopsis thaliana* | thale cress | 735 |
| *M. musculus* | house mouse | 722 |
| *Acropora digitifera* | Acropora digitifera | 710 |
| *Trichuris suis* | pig whipworm | 679 |
| *Crassostrea gigas* | Pacific oyster | 668 |

To understand global gene content evolution in the *P. waltl* proteome, we generated clusters of orthologous and paralogous gene families comparing the *P. waltl* proteome with those of 9 other vertebrates (Table 4). The OrthoFinder program identified 18,559 orthogroups consisting of 215,304 genes. The *P. waltl* proteome was clustered into 15,923 orthogroups, among which 13,283 and 14,183 groups were shared with human and *X. laevis*, respectively (Table 4; Fig. 2). We found 660 orthologous groups, consisting of 2,958 genes, that are unique to *P. waltl*; these loci presumably represent evolutionarily young genes or loci that have undergone considerable divergence following gene duplications. These lineage-specific genes might account for the traits unique to *P. waltl.* Additionally, we found 784 orthologous groups that are shared only among amphibians (*P. waltl*, and *X. tropicalis*).

**Figure 2.**
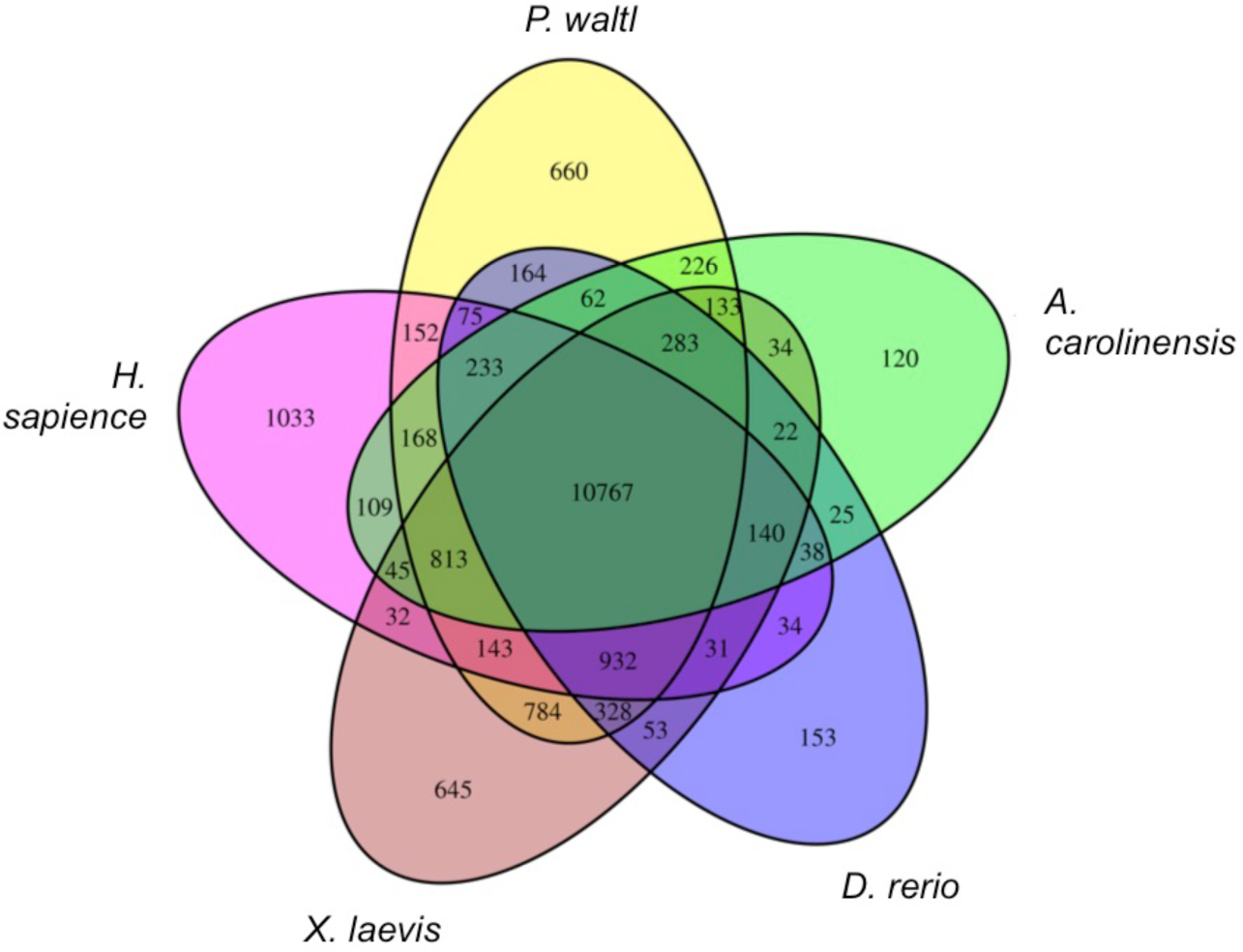
Venn diagram of shared and unique orthogroups in five vertebrates. Orthogroups were identified by clustering of orthologous groups using OrthoFinder.

**Table 4.**
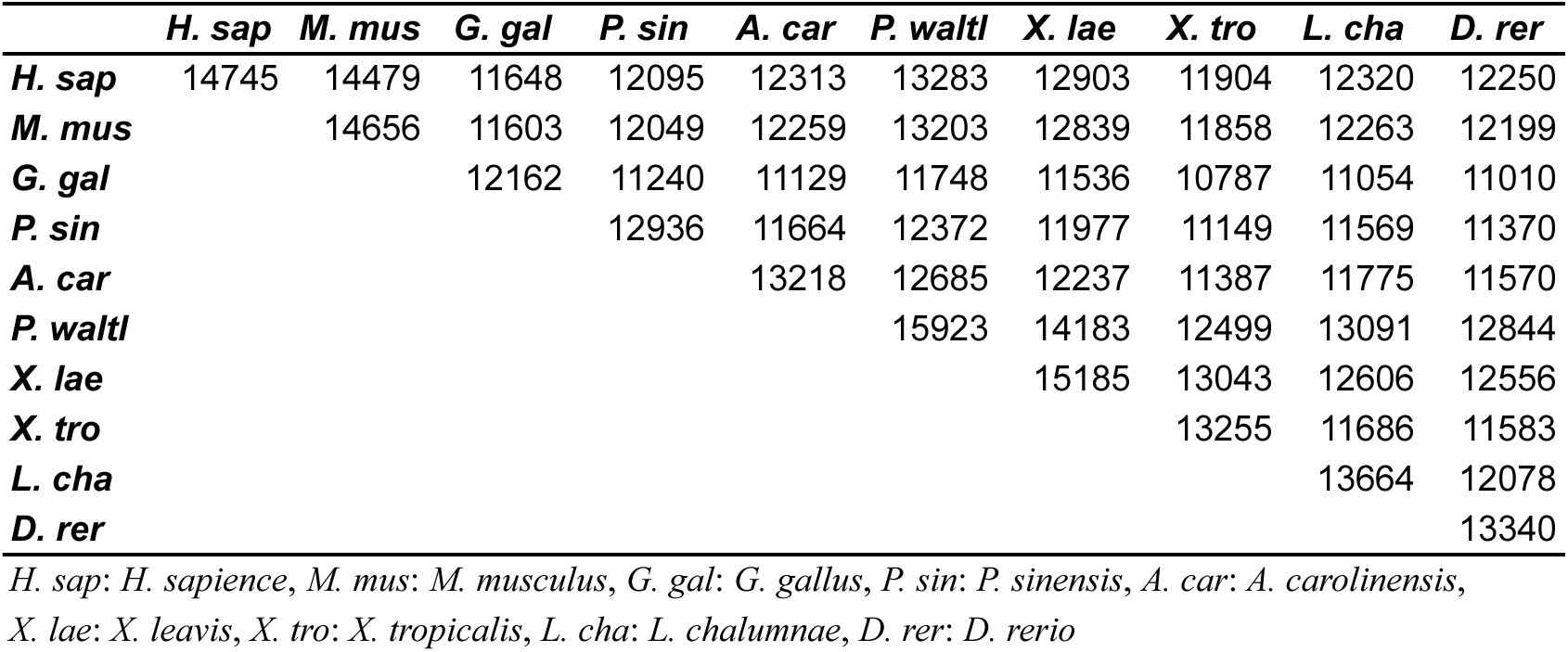
Orthogroup overlaps.

Salamanders are another group of urodele amphibians. We compared our *P. waltl* transcriptome with that of axolotl and identified 22,907 orthologous groups from the pairwise comparison. These two species shared 22,307 orthologous groups, while retaining 321 and 279 species-specific groups, respectively (Supplementary figure 1). In both organisms, these species-specific groups often contained LINE elements. Previous reports have shown that LINE elements are abundant in urodele amphibian genomes^24,26^. Thus, we speculated that genes containing LINE elements have evolved more rapidly, accumulating lineage-specific mutations as a result of retrotransposition events. These LINE elements might be related to species-specific regeneration abilities, given that LINE elements are known to be activated during salamander limb regeneration^62^, although the functional contribution of these loci in regeneration remains hypothetical.

### Gene co-expression pattern analysis

We quantified gene expression and profiled the expression patterns across all of the samples examined. A multi-dimensional scaling (MDS) plot of the 29 samples was used to depict the transcriptome similarities among the samples (Fig. 3). Samples derived from differentiated tissues/organs of adults (red dots in Fig. 3) yielded transcriptomes that were clearly distinct from those of samples from developing embryos (blue dots in Fig. 3). Samples at similar developmental stages clustered closer to each other than to those of differentiated tissues/organs; samples derived from the amputation experiments clustered on the MDS plot based on the amputated tissue. Notably, directional distances on the dimension-2 axis indicated a continuum in the direction of changes that was consistent with developmental progression. Specifically, the embryonic samples were clearly ordered along the dimension-2 axis from unfertilized egg to gastrula to neurula to tail-bud stage, implying that gene expression gradually changes with progression during embryogenesis.

**Figure 3.**
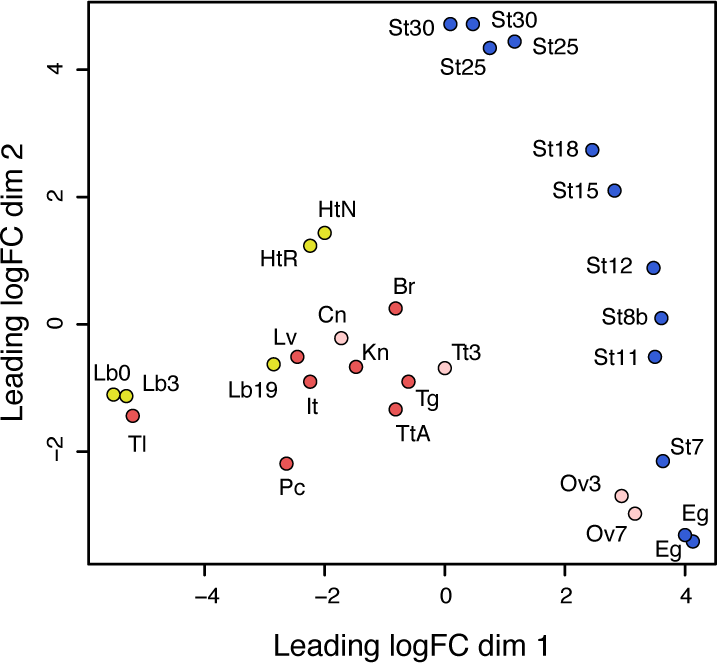
MDS plot for RNA-Seq gene expression of *P. waltl* tissues, organs, and embryogenesis samples. Multi-dimensional scaling (MDS) plot showing relatedness between transcript expression profiles of organs, tissues, and embryos of *P. waltl* at different developmental stages. Red dots represent the expression profiles of adult tissues/organs and pink dots represent those of juveniles (3 or 7 months). The labels indicate the tissues and sources as follows, Br: brain (adult), Cn: connective tissue (3 months), It: intestine (adult), Kn: kidney (adult), Lv: liver (adult), Ov3: ovary (3 months), Ov7: ovary (7 months), Pc: pancreas (adult), Tg: testicular gland (adult), TtA: testis (adult), and Tt3: testis (3 months). Blue dots represent the expression profiles during embryogenesis. The labels indicate the stages as follows, Eg: unfertilized egg, St7: stage 7 (late blastula), St8b: stage 8b (early gastrula), St11: stage 11 (middle gastrula), St12: stage 12 (late gastrula), St15: stage 15 (neural plate stage), St18: stage 18 (late neural fold stage), St25: stage 25 (tail-bud stage), and St30: stage 30 (gill protrusion stage). Yellow dots represent the expression profiles in the regeneration process after amputation, where the labels Lb0, Lb3, and Lb19 indicate limb or limb blastema expression profiles at 0, 3, and 19 dpa (respectively); HtR and HtN indicate expression profiles of the hearts regenerating after amputation and in unamputated controls (respectively).

To understand the co-expression relationships between genes at a systems level, we performed WGCNA. This unsupervised and unbiased analysis identified distinct co-expression modules corresponding to clusters of correlated transcripts (Fig. 4). WGCNA identified 21 co-expressed modules from the expression data spanning 29 samples; each module contained 53 to 3283 co-expressed genes (Fig. 5). Each module represents genes with highly correlated expression profiles, either in a single tissue or in a narrow window of developmental stages. Out of 21 modules, 11 represent a tissue-specific pattern in the adult tissues: the modules indicated by different colors represent different tissues (blue, brown, red, black, pink, green, yellow, cyan, light green, green, grey, and royal blue for testis, ovary, intestine, pancreas, liver, testicular gland, brain, tail, heart, testicular connective tissue, and kidney-specific expression patterns, respectively) (Fig. 5A). Six modules were embryonic (Fig. 5B). The turquoise module was composed of 3283 genes whose expression was observed only in unfertilized eggs, representing maternal transcripts that functioned in the early stages during *P. waltl* embryogenesis. On the other hand, modules indicated in purple, yellow, dark red, midnight blue, and light yellow to genes exhibiting zygotic co-expression after the mid-blastula transition (MBT; St 6-7), with the modules representing a progressive pattern showing peaks at St 8b-12, St 15, St 18, St 25-30 and St 30, respectively. In the regeneration experiments, limb-enriched genes were clustered into three modules based on a pattern corresponding to the responsiveness to amputation treatment: genes designated in salmon, magenta, and light cyan showed peaks at 0, 3, and 19 days post amputation (dpa) (Fig. 5C).

**Figure 4.**
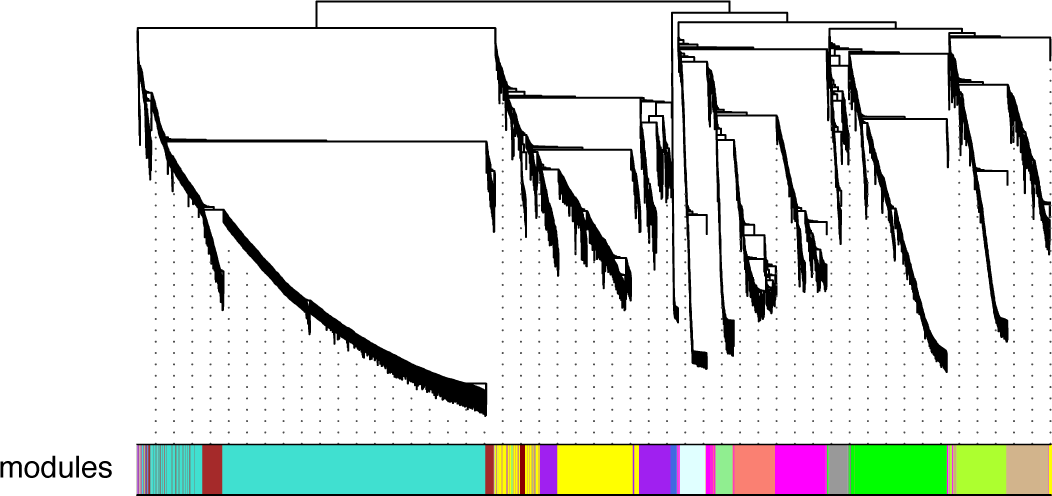
Gene co-expression analysis of *P. waltl* transcriptome. Hierarchical cluster tree of the *P. waltl* genes showing co-expression modules identified using WGCNA. Modules correspond to branches and are labelled by colors as indicated by the color band underneath the tree.

**Figure 5.**
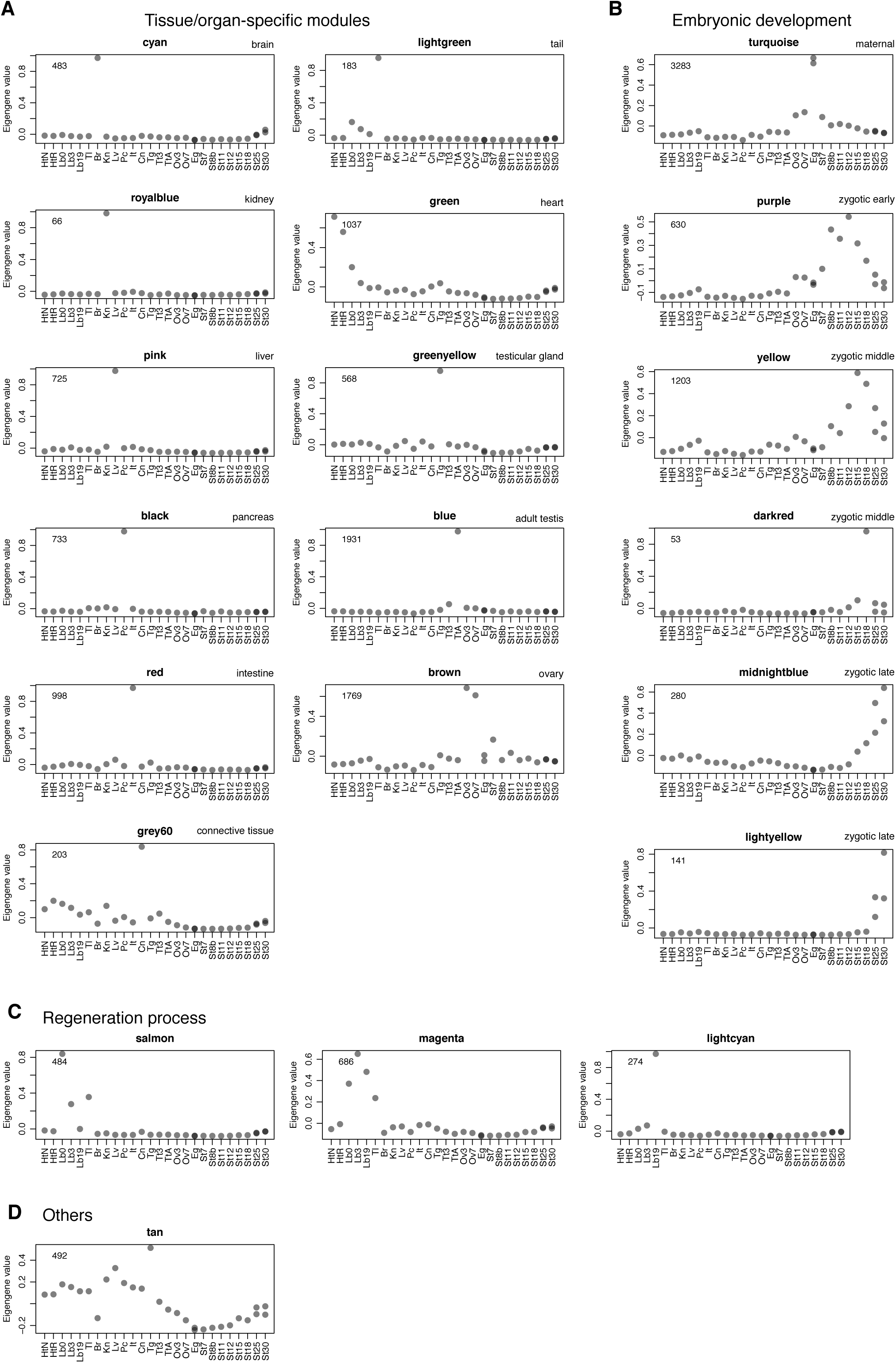
Co-expression gene modules. The co-expression gene modules identified using WGCNA are shown. Each grey dot represents the value of the respective module’s Eigengene. The number at the top left in each panel indicates the number of genes belonging to a module exhibiting unique expression. The modules are classified into 4 categories based on the expression pattern: modules associated with (A) specific tissues/organs, (B) embryogenesis, (C) regeneration processes, and (D) others. The sample abbreviations indicated by labels at the bottom of each panel are defined in the Figure 3 legend.

### Major signaling pathways

Cell signaling pathways are essential for embryogenesis and organogenesis and are highly conserved in vertebrates. We inspected the gene repertoire of major signal-factor encoding genes and analyzed these expression patterns at the various developmental stages (Supplementary figure 2). It turned out that the repertoire of signaling genes of *P. waltl* is typical for vertebrates, but we found a few cases of urodele amphibian-specific gene losses and duplications. An example is bmp gene family. Orthologs of *bmp2*, *bmp7*, and *bmp16* but not of *bmp4*, were identified in the transcriptome of *P. waltl.* Furthermore, no orthologs of *bmp4* were identified in the transcriptomes of two other urodeles, *A. mexicanum*, and *C. pyrrhogaster* (Fig. 6). Although *bmp16* has been thought to be confined to only teleost fish species^63^, we found urodele *bmp16* with accompanying ortholog of reptile *A. carolinesis.* Thus, our phylogenetic analysis suggested that urodeles and anurans have lost *bmp4* and *bmp16*, respectively, in each lineage. In the anuran *Xenopus* species, *bmp2* is maternally expressed, and *bmp4* and *bmp7* are zygotically expressed^55,64^. *bmp7* and *bmp2* showed high- and low-level maternal expression (respectively) in *P. waltl* (Supplementary figure 2D), suggesting that the functions of *bmp2*, *bmp4*, and *bmp7* are redundant in amphibian species. Expression of *P. waltl bmp16* was not apparent in early developmental stages (Supplementary figure 2D). The *wnt* gene family set is conserved in *P. waltl* as in other vertebrates. But we detected two additional paralogous genes encoding Wnt ligands, *wnt11b* and *wnt7*-*like* (Supplementary figure 2A); we postulate that these additional loci were generated by duplication in the lineage leading to *P. waltl*. In vertebrates, six highly conserved *igfbp*-family genes are typically observed, and *P. waltl* has all six *igfbp* orthologous genes (Supplementary figure 2E), while *Xenopus* lacks *igfbp3* and *igfbp6* orthologs^65^.

**Figure 6.**
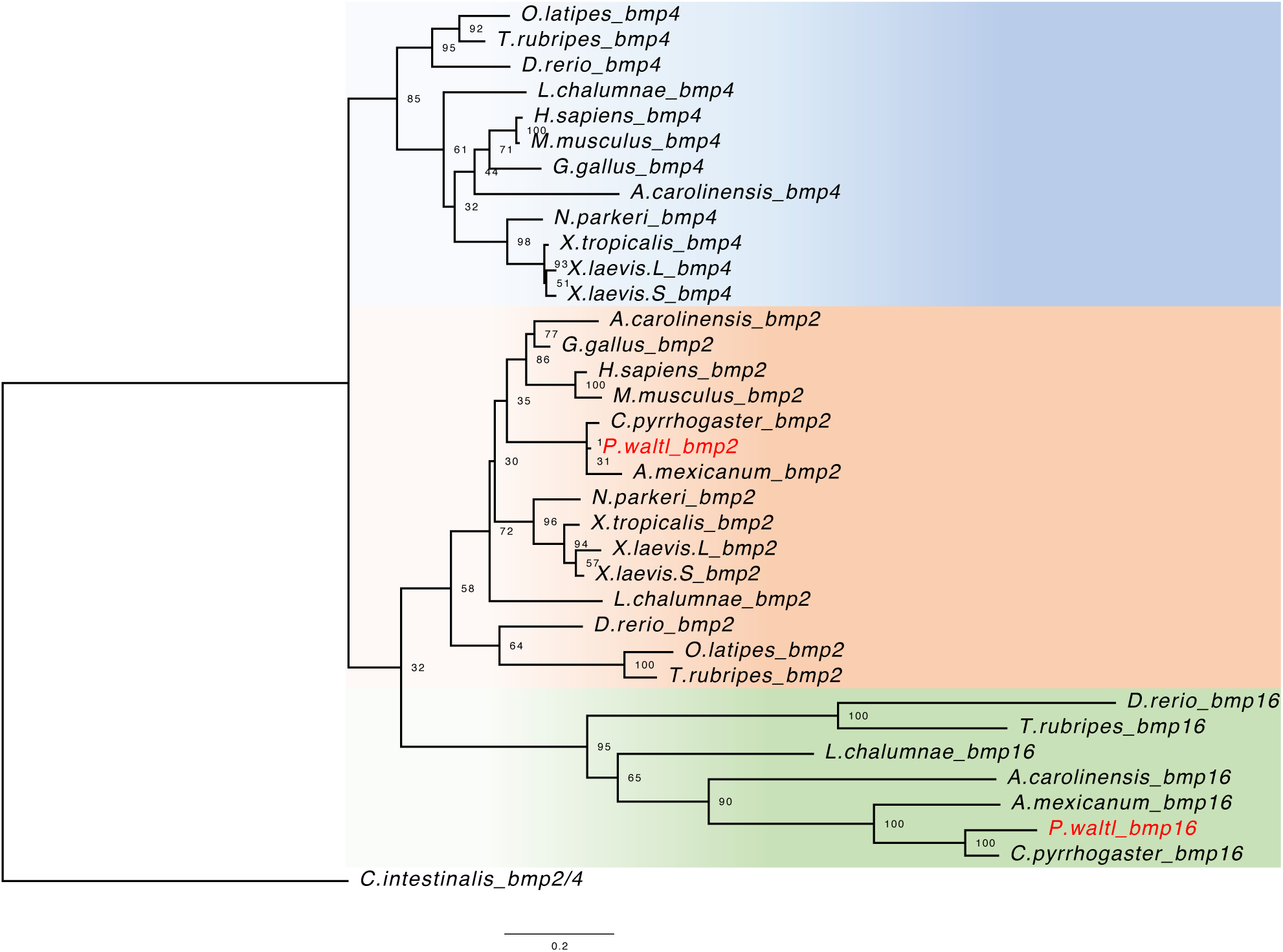
Phylogenetic tree of *bmp2/4/16* genes among vertebrates. The phylogenetic tree was reconstructed using 34 vertebrate orthologs, including 12 *bmp4*, 15 *bmp2*, and 7 *bmp16* genes; an ascidian *bmp2/4* was used as the outgroup. The number at each node represents the bootstrap probability.

In sum, most of the orthologous genes for major signal molecules were identified in *P. waltl*, which therefore harbors a gene repertoire typical of vertebrates, with a few exceptions. The expression patterns of the signaling molecule-encoding genes of *P. waltl* sometimes differed from those of *Xenopus* species. Further research on these differences is expected to expand our understanding of the evolution and development of amphibians.

### *Hox* genes and their expression dynamics during embryogenesis

In tetrapod genomes, approximately 40 *hox* genes are present and organized into four *hox* clusters. In amphibians, the genes encoding Hoxb13 and Hoxd12 have been lost, while the Hoxc3-encoding gene is retained^66^. No *hoxc1* ortholog has been identified in amphibians, with the exception of caecillians^67^. The genome of the diploid *X. tropicalis* and the allotetraploid *X. laevis* harbors 38 and 75 functional *hox* genes, respectively^68^. Thus, usual amphibians appear to have retained 38 *hox* genes per diploid genome. Consistent with this observation, we identified a complete set of all of the *hox* gene orthologs in the *P. waltl* transcriptome (Fig. 7).

**Figure 7.**
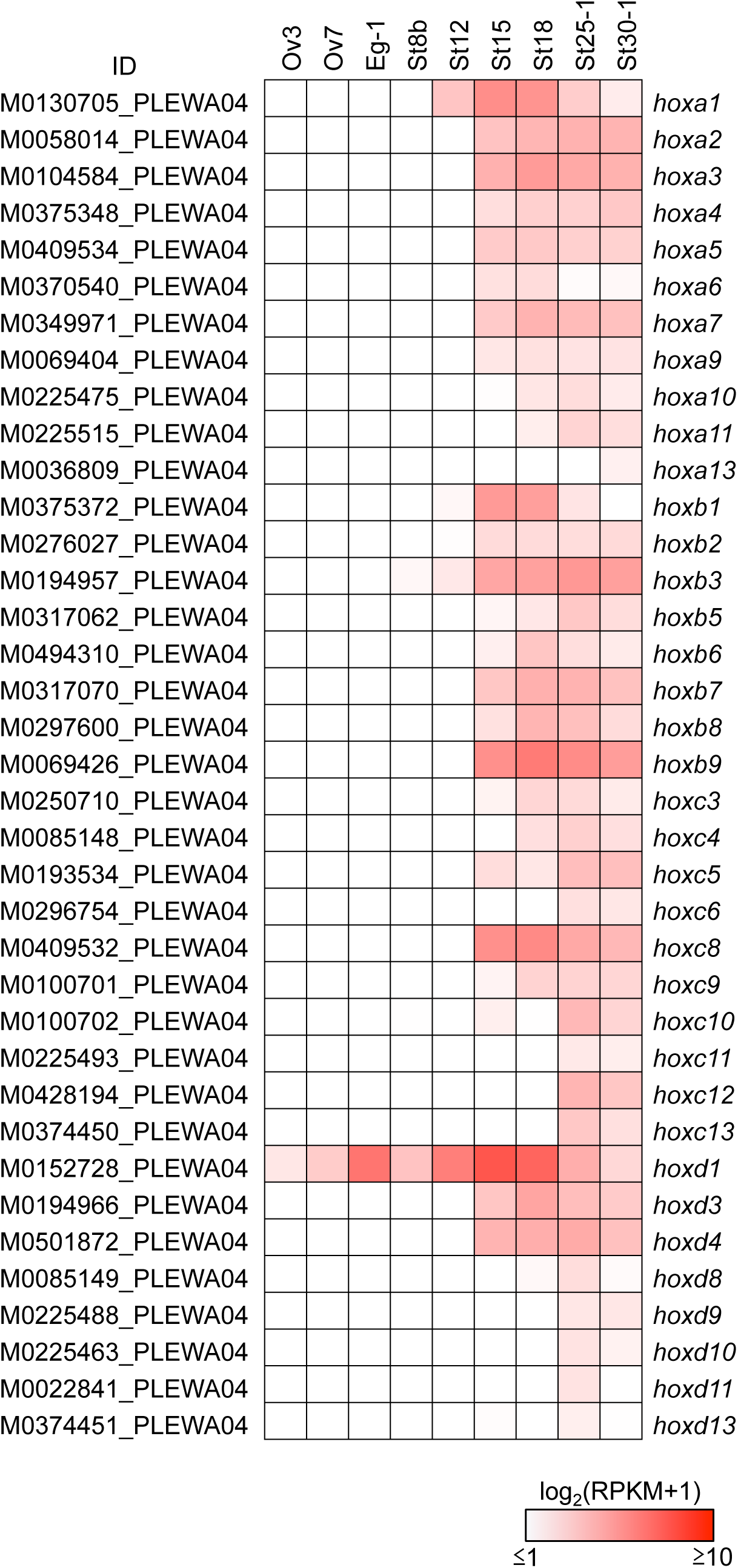
Expression profile of *hox* genes during oogenesis and embryogenesis. A total of 37 *hox* genes are listed from the assembly data of PLEWA04. The sample abbreviations indicated by labels at the bottom of each panel are t in the Figure 3 legend. Ovaries were sampled at three and six months after metamorphosis (Ov3 and Ov6, respectively). Note that *P. waltl hoxb13*, *hoxcl*, and *hoxd12* orthologs were not identified from our transcriptome data. Most of the *hox* genes were zygotically activated; only the *hoxd1* mRNA was synthesized through oogenesis and stored at the one-cell stage. RPKM values of each gene are indicated as a color gradient on a log_10_ scale, ranging from red (maximum) to white (minimum).

The expression profile of the *P. waltl hox* genes of during embryogenesis was similar to those of axolotl and *Xenopus*, suggesting that the regulation of this gene family is conserved among amphibians (Fig. 7)^66,68^. Anterior *hox* genes were activated starting around the time of the MBT; posterior *hox* genes were gradually up-regulated at the late embryonic stage, reflecting their spatio-temporal collinearity during embryogenesis. Interestingly, *hoxd1* of *P. waltl* was found to be stored as a maternal mRNA at the oocyte and one-cell stage (Fig. 7), whereas the orthologous genes were expressed after MBT in axolotl and *Xenopus*^66,68,69^.

The correlation between newt genomic gigantism and remarkable regenerative ability has been interpreted to suggest that the genome of a prototypical newt underwent species-specific whole genome duplication. Because *hox* genes are maintained as highly conserved gene clusters in vertebrate genomes, the number of *hox* clusters usually reflects the number of whole genome duplications each genome experienced during evolution^70^. In our *P. waltl* gene repertoire, we only found one-to-one orthologs of *hox* genes when comparing among available amphibian genes. This result suggested that the newt genome did not undergo additional whole genome duplication. Similarly, the recently published axolotl giant genome showed no evidence for additional whole genome duplication; instead, the axolotl genome has a correspondingly enlarged genic component, primarily due to the presence of especially long introns^24,26,71^. In the salamander genome, expansion of LTR retrotransposons also contributes to genome gigantism^72^. Such mechanisms also may have contributed to newt genome gigantism and may be related to the incredible regenerative ability of this species.

### Transcriptomic features of regenerating limbs

Numerous developmental pathway genes have been reported to be reactivated during limb regeneration. For example, *hox13*-paralogous genes are expressed in distal regions of developing and regenerating amphibian limbs^73-76^. Indeed, the present work confirmed the reactivation of the *hoxa/c/d13* genes in regenerating *P. waltl* forelimb at 19 dpa, when the blastema is fully formed (Fig. 8A). Other transcription factor-encoding genes known to be involved in limb development and regeneration (*msx1*, *msx2*, *prrx1*, *prrx2*, *tbx5*, and *hand2*) also were drastically up-regulated at 19 dpa, indicating that the transcriptome defined here successfully recaptured the expression pattern of limb regeneration (Fig. 8A). Consistent with the reports in axolotl^57^, genes encoding RNA-binding proteins (*rbmx*, *hnrnpa1*, and *sfrs1*), matrix metalloproteinases (*mmp3* and *mmp11*), and a novel proteinase inhibitor (*kazald1*) were up-regulated in regenerating limb at 19 dpa in *P. waltl* (Supplementary figure 3). These results suggested that these genes are commonly involved in regeneration in the two urodele amphibians.

**Figure 8.**
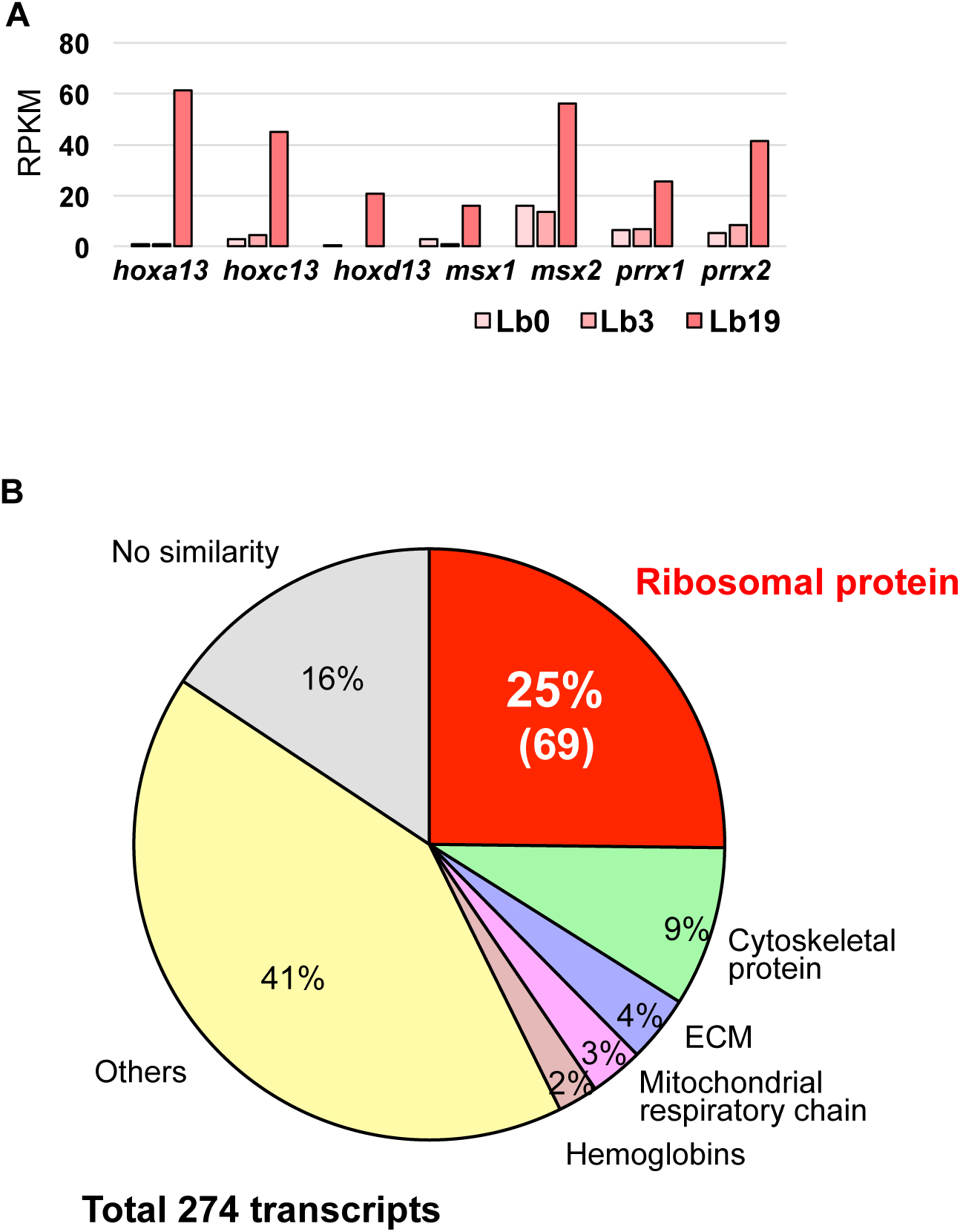
Expression profile of regenerating limb-enriched genes. (a) Expression of transcription factor-encoding genes involved in limb development during regeneration. The *hox13*, *msx1* and *2*, *prrx1* and *2*, *tbx5*, and *hand2* genes were significantly up-regulated in the forelimb at 19 dpa. RPKM values of each gene were determined from the assembly data of PLEWA04. (b) Details of co-expressed genes in regenerating limb at 19 dpa. A total of 274 genes in this WGCNA module (indicated by light cyan symbols in Fig. 5) were identified. Notably, genes encoding proteins of the large and small ribosomal subunits accounted for 25% (69 out of 274) of the genes in this module.

Intriguingly, WGCNA revealed a unique transcriptomic feature of regeneration (Fig. 5; light cyan symbols). The light cyan module contained 274 genes that are co-regulated in regenerating limb at 19 dpa (Fig. 8B). Notably, this module included 69 ribosomal protein-encoding genes (Fig. 8B). These transcripts typically were not detected in other tissues and organs, suggesting that these ribosomal protein-encoding genes are likely to have be limb- or regeneration-specific roles. These ribosomal proteins may contribute to organ remodeling *via* regeneration-specific protein synthesis. Consistent with this inference, the expression of ribosomal proteins is down-regulated in *Xenopus* spinal cord regeneration at the non-regenerative stage after metamorphosis^77^.

Axolotl is another good model organism of regeneration; that organism also is a urodele amphibian, and the axolotl genome was recently reported^26^. Axolotl is a neotenic animal, that is, one that retains aspects of the larval state even after sexual maturation^12^. Interestingly, axolotl shows restricted regenerative capacity compared to newts^24^. How did such differences in metamorphosis and regenerative capacity arise despite the closely related nature of these species? Our near-complete *P. waltl* gene catalogue, together with the recently reported *P. waltl* draft genome sequence^24^, is expected to facilitate genome-wide comparisons between these two model urodele amphibians.

## Conclusions

In the present study, we built a reference gene catalogue of *P. waltl* using transcriptome data sets generated from a wide variety of samples. As a BUSCO analysis showed, our gene models appear to cover most of the protein coding genes on the newt genome. The near-complete gene catalogue and the associated information will be valuable resources for any researchers to use *P. waltl.* To share the resources in the community, we established a portal website, designated iNewt (http://www.nibb.ac.jp/imori/main/), where these transcriptome data such as gene models, annotations and expression profiles can be obtained. The portal site also permits BLAST searches against the data set. With these references, *P. waltl* is promising to serve a good model to expand our understanding of molecular mechanisms underlying regeneration. Given the newts’ unique biological properties, we further expect that our reference gene catalogue, together with the technique of highly efficient CRISPR/Cas9 genome editing^41^, will open new avenues for researches using *P. waltl* besides regeneration which includes developmental biology, stem cell biology, cancer research, reproductive biology, evolutionary biology, ethology, and toxico-genomics.

## Acknowledgments

This work was supported by MEXT/JSPS KAKENHI (Grant Numbers JP16H01254 TH, JP16K08467 TH, JP17J04796 MS, JP16K18613 MM, JP17K14980 YH, JP16H04794 TT, JP15K06802 KTS) and Grant for Basic Science Research Projects of the Sumitomo Foundation to MM (No. 170845), and Chuo University Personal Research Grant to AF. This work was also supported by NIBB Collaborative Research Program (17-431, 18-204) and computations were partially performed on the NIG supercomputer at ROIS. Kyorin Corporation (Hyogo, Japan) kindly provided the feeds for the newts.

## References

1. Spemann, H., and Mangold, O. 1924, Über Induktion von Embryonalanlagen durch Implantation artfremder Organisatoren, Arch. Mikrosk. Anat. En., 100, 599–638

2. Wolff, G. 1895, Entwicklungsphysiologische Studien 1. Die regeneration der Urodelenlinse, Wilhelm Roux’s Arch. Entwined Meghan Org., 1, 380-390.

3. Eguchi, G., Abe, S. I., and Watanabe, K. 1974, Differentiation of lens-like structures from newt iris epithelial cells in vitro, Proc. Natl. Acad. Sci. U S A, 71, 5052-5056.

4. Agata, K., and Inoue, T. 2012, Survey of the differences between regenerative and non-regenerative animals, Dev Growth Differ 54, 143–152.

5. Hayashi, T., Mizuno, N., and Kondoh, H. 2008, Determinative roles of FGF and Wnt signals in iris-derived lens regeneration in newt eye, Dev. Growth. Differ., 50, 279-287.

6. Inoue, T., Inoue, R., Tsutsumi, R., Tada, K., Urata, Y., Michibayashi, C. et al. 2012, Lens regenerates by means of similar processes and timeline in adults and larvae of the newt *Cynops pyrrhogaster*, Dev. Dyn., 241, 1575-1583.

7. Maki, N., Takechi, K., Sano, S., Tarui, H., Sasai, Y., and Agata, K. 2007, Rapid accumulation of nucleostemin in nucleolus during newt regeneration, Dev. Dyn., 236, 941-950.

8. Maki, N., Tsonis, P. A., and Agata, K. 2010, Changes in global histone modifications during dedifferentiation in newt lens regeneration, Mol. Vis., 16, 1893-1897.

9. Brockes, J. P. 1997, Amphibian limb regeneration: rebuilding a complex structure’, Science, 276, 81-87.

10. Tsutsumi, R., Inoue, T., Yamada, S.,and Agata, K. 2015, Reintegration of the regenerated and the remaining tissues during joint regeneration in the newt *Cynops pyrrhogaster*, Regeneration (Oxf), 2, 26-36.

11. Mercer, S. E., Odelberg, S. J., and Simon, H. G. 2013, A dynamic spatiotemporal extracellular matrix facilitates epicardial-mediated vertebrate heart regeneration, Dev. Biol., 382, 457-469.

12. Kurosaka, H., Takano-Yamamoto, T., Yamashiro, T., and Agata, K. 2008, Comparison of molecular and cellular events during lower jaw regeneration of newt (*Cynops pyrrhogaster*) and West African clawed frog (*Xenopus tropicalis*), Dev. Dyn., 237, 354-365.

13. Ikegami, Y., Mitsuda, S., and Araki, M. 2002, Neural cell differentiation from retinal pigment epithelial cells of the newt: an organ culture model for the urodele retinal regeneration, J. Neurobiol., 50, 209-220.

14. Chiba, C., Oi, H., and Saito, T. 2005, Changes in somatic sodium currents of ganglion cells during retinal regeneration in the adult newt, Brain Res. Dev. Brain Res., 154, 25-34.

15. Okamoto, M., Ohsawa, H., Hayashi, T., Owaribe, K., and Tsonis, P. A. 2007, Regeneration of retinotectal projections after optic tectum removal in adult newts, Mol. Vis., 13, 2112-2118.

16. Berg, D. A., Kirkham, M., Beljajeva, A., Knapp, D., Habermann, B., Ryge, J. et al. 2010, Efficient regeneration by activation of neurogenesis in homeostatically quiescent regions of the adult vertebrate brain, Development, 137, 4127-4134.

17. Urata, Y., Yamashita, W., Inoue, T., and Agata, K. 2018, Spatio-temporal neural stem cell behavior leads to both perfect and imperfect structural brain regeneration in adult newts, Biol. Open, 7 (6).

18. Zhang, F., Clarke, J. D., Santos-Ruiz, L., and Ferretti, P. 2002, Differential regulation of fibroblast growth factor receptors in the regenerating amphibian spinal cord in vivo, Neuroscience, 114, 837-848.

19. Grubb, R. B. 1975, An autoradiographic study of the origin of intestinal blastemal cells in the newt, *Notophthalmus viridescens*, Dev. Biol., 47, 185-195.

20. Uchida, T., and Hanaoka, K. I., 1949, The occurrence of oviform cells by hormonal injection in the regenerated testes of a newt, Cytologia, 15, 109–130.

21. Flament, S., Dumond, H., Chardard, D., and Chesnel, A. 2009, Lifelong testicular differentiation in *Pleurodeles waltl* (Amphibia, Caudata), Reprod. Biol. Endocrinol., 7, 21.

22. Romanova, 1959, Regenerating potential of a hypertrophied lung in *Triturus cristatus*, Biull. Eksp. Biol. Med., 47, 89-94.

23. Tsutsumi, R., Yamada, S., and Agata, K. 2016, Functional joint regeneration is achieved using reintegration mechanism in Xenopus laevis, Regeneration (Oxf), 3, 26-38.

24. Elewa, A., Wang, H., Talavera-López, C., Joven, A., Brito, G., Kumar, A., et al. 2017. Reading and editing the *Pleurodeles waltl* genome reveals novel features of tetrapod regeneration. Nat. Commun., 8, 2286.

25. Gregory, T. R. 2005, Synergy between sequence and size in large-scale genomics, Nat. Rev. Genet., 6, 699-708.

26. Nowoshilow, S., Schloissnig, S., Fei, J.F., Dahl, A., Pang, A.W.C., Pippel, M., et al. 2018, The axolotl genome and the evolution of key tissue formation regulators, Nature 554, 50-55.

27. Ingram, A. J. 1972, The lethal and hepatocarcinogenic effects of dimethylnitrosamine injection in the newt *Triturus helveticus*, Br. J. Cancer, 26, 206-215.

28. Okamoto, M. 1997, Simultaneous demonstration of lens regeneration from dorsal iris and tumour production from ventral iris in the same newt eye after carcinogen administration, Differentiation, 61, 285-92.

29. Ueno, T., Ohgami, T., Harada, Y., Ueno, S., Iwao, Y. 2014, Egg activation in physiologically polyspermic newt eggs: involvement of IP_3_ receptor, PLC*γ*, and microtubules in calcium wave induction, Int. J. Dev. Biol., 58, 315-323.

30. Uribe, M.C., and Mejía-Roa, V. 2014, Testicular structure and germ cells morphology in salamanders. Spermatogenesis 4, e988090.

31. Kikuyama, S., Toyoda, F., Ohmiya, Y., Matsuda, K., Tanaka, S., and Hayashi, H. 1995, Sodefrin: a female-attracting peptide pheromone in newt cloacal glands, Science, 267 (5204), 1643-1645.

32. Nakada, T., Toyoda, F., Matsuda, K., Nakakura, T., Hasunuma, I., Yamamoto, K., et al. 2017, Imorin: a sexual attractiveness pheromone in female red-bellied newts (*Cynops pyrrhogaster*), Sci. Rep., 7, 41334.

33. Mouchet, F., Gauthier, L., Mailhes, C., Ferrier, V., and Devaux, A. 2006, Comparative evaluation of genotoxicity of captan in amphibian larvae (*Xenopus laevis* and *Pleurodeles waltl*) using the comet assay and the micronucleus test, Environ. Toxicol. 21, 264-277.

34. Mouchet, F., Gauthier, L., Baudrimont, M., Gonzalez, P., Mailhes, C., Ferrier, V., et al. 2007, ‘Comparative evaluation of the toxicity and genotoxicity of cadmium in amphibian larvae (*Xenopus laevis* and *Pleurodeles waltl*) using the comet assay and the micronucleus test’, Environ. Toxicol., 22, 422-435.

35. Hirako, A., Takeoka, Y., Hayashi, T., Takeuchi, T., Furukawa, S., and Sugiyama, A. 2017, Effects of cadmium exposure on Iberian ribbed newt, J. Toxicol. Pathol. 30, 345-350.

36. Bour, A., Mouchet, F., Cadarsi, S., Silvestre, J., Verneuil, L., Baqué, D. et al. 2016, Toxicity of CeO_2_ nanoparticles on a freshwater experimental trophic chain: A study in environmentally relevant conditions through the use of mesocosms, Nanotoxicology, 10, 245-255.

37. Makita, R., Kondoh, H., and Okamoto, M. 1995, Transgenesis of newt with exogenous gene expression facilitated by satellite 2 repeats, Dev. Growth and Differ., 37, 605-616.

38. Ueda, Y., Kondoh, H., and Mizuno, N. 2005, Generation of transgenic newt *Cynops pyrrhogaster* for regeneration study, Genesis 41, 87-98.

39. Hayashi, T., Yokotani, N., Tane, S., Matsumoto, A., Myouga, A., Okamoto, M. et al. 2013, Molecular genetic system for regenerative studies using newts, Dev. Growth Differ. 55, 229-236.

40. Hayashi, T., Sakamoto, K., Sakuma, T., Yokotani, N., Inoue, T., Kawaguchi, E. et al. 2014, Transcription activator-like effector nucleases efficiently disrupt the target gene in Iberian ribbed newts (*Pleurodeles waltl*), an experimental model animal for regeneration, Dev. Growth Differ. 56, 115-121.

41. Suzuki, M., Hayashi, T., Inoue, T., Agata, T., Hirayama, T., Suzuki, T. et al. 2018, Cas9 ribonucleoprotein complex allows direct and rapid analysis of coding and noncoding regions of target genes in *Pleurodeles waltl* development and regeneration. Dev. Biol., in press.

42. Shi, D.L. and Boucaut, J.C. 1995, The chronological development of the urodele amphibian *Pleurodeles waltl* (Michah). Int. J. Dev. Biol. 39, 427-441.

43. Grabherr, M. G., Haas, B. J., Yassour, M., Levin, J. Z., Thompson, D. A., Amit, I. et al. 2011, Full-length transcriptome assembly from RNA-Seq data without a reference genome, Nat. Biotechnol., 29, 644-652.

44. Bolger, A.M., Lohse, M., and Usadel, B. 2014, Trimmomatic: a flexible trimmer for Illumina sequence data, Bioinformatics, 30, 2114-2120.

45. Haas, B. J., Papanicolaou, A., Yassour, M., Grabherr, M., Blood, P. D., Bowden, J. et al. 2013, De novo transcript sequence reconstruction from RNA-seq using the Trinity platform for reference generation and analysis, Nat. Protoc., 8, 1494-1512.

46. Fu, L., Niu, B., Zhu, Z., Wu, S., and Li, W. 2012, CD-HIT: accelerated for clustering the next-generation sequencing data, Bioinformatics, 28, 3150-3152.

47. Simão, F.A., Waterhouse, R.M., Ioannidis, P., Kriventseva, E.V., and Zdobnov, E.M. 2015, BUSCO: assessing genome assembly and annotation completeness with single-copy orthologs, Bioinformatics, 31, 3210-3212.

48. Hara, Y., Tatsumi, K., Yoshida, M., Kajikawa, E., Kiyonari, H., and Kuraku, S. 2015, Optimizing and benchmarking de novo transcriptome sequencing: from library preparation to assembly evaluation. BMC Genomics, 16, 977.

49. Götz, S., García-Gómez, J.M., Terol, J., Williams, T.D., Nagaraj, S.H., Nueda, M.J. et al. 2008, High-throughput functional annotation and data mining with the Blast2GO suite. Nucleic Acids Res., 36, 3420-3435.

50. Emms, D.M. and Kelly, S. 2015, OrthoFinder: solving fundamental biases in whole genome comparisons dramatically improves orthogroup inference accuracy, Genome Biol., 16, 157.

51. Bray, N.L., Pimentel, H., Melsted, P., and Pachter, L. 2016, Near-optimal probabilistic RNA-seq quantification, Nat. Biotechnol., 34, 525-527.

52. Robinson, M.D., McCarthy, D.J., and Smyth, G.K. 2010, edgeR: a Bioconductor package for differential expression analysis of digital gene expression data, Bioinformatics, 26, 139-140.

53. Langfelder, P. and Horvath, S. 2008, WGCNA: an R package for weighted correlation network analysis. BMC Bioinformatics, 9, 559.

54. Zerbino, D. R., Achuthan, P., Akanni, W., Amode, M. R., Barrell, D., Bhai, J. et al. 2018, Ensembl 2018, Nucleic Acids Res., 46 (D1), D754-D61.

55. Session, A. M., Uno, Y., Kwon, T., Chapman, J. A., Toyoda, A., Takahashi, S. et al. 2016, Genome evolution in the allotetraploid frog Xenopus laevis, Nature, 538, 336-343.

56. Sun, Y. B., Xiong, Z. J., Xiang, X. Y., Liu, S. P., Zhou, W. W., Tu, X. L. et al. 2015, Whole-genome sequence of the Tibetan frog *Nanorana parkeri* and the comparative evolution of tetrapod genomes, Proc. Natl. Acad. Sci. U S A, 112, E1257-62.

57. Bryant, D.M., Johnson, K., DiTommaso, T., Tickle, T., Couger, M.B., Payzin-Dogru, D., Lee, T.J., Leigh, N.D., Kuo, T.H., Davis, F.G., et al. (2017). A Tissue-Mapped Axolotl De Novo Transcriptome Enables Identification of Limb Regeneration Factors. Cell Rep 18, 762-776.

58. Nakamura, K., Islam, M. R., Takayanagi, M., Yasumuro, H., Inami, W., Kunahong, A. et al. 2014, A transcriptome for the study of early processes of retinal regeneration in the adult newt, Cynops pyrrhogaster, PLoS One, 9, e109831.

59. Edgar, R. C. 2004, MUSCLE: multiple sequence alignment with high accuracy and high throughput, Nucleic Acids Res., 32, 1792-1797.

60. Kumar, S., Stecher, G., and Tamura, K. 2016, MEGA7: Molecular Evolutionary Genetics Analysis Version 7.0 for Bigger Datasets, Mol. Biol. Evol., 33, 1870-1874.

61. Irisarri, I., Baurain, D., Brinkmann, H., Delsuc, F., Sire, J. Y., Kupfer, A. et al. 2017, Phylotranscriptomic consolidation of the jawed vertebrate timetree, Nat. Ecol. Evol. 1, 1370-1378.

62. Zhu, W., Kuo, D., Nathanson, J., Satoh, A., Pao, G. M., Yeo, G. W. et al. 2012, Retrotransposon long interspersed nucleotide element-1 (LINE-1) is activated during salamander limb regeneration, Dev. Growth Differ., 54, 673-685.

63. Feiner, N., Begemann, G., Renz, A. J., Meyer, A., Kuraku, S. 2009, The origin of bmp16, a novel Bmp2/4 relative, retained in teleost fish genomes, BMC. Evol. Biol., 9, 277.

64. Owens, N. D. L., Blitz, I. L., Lane, M. A., Patrushev, I., Overton, J. D., Gilchrist, M. J. et al. 2016, Measuring Absolute RNA Copy Numbers at High Temporal Resolution Reveals Transcriptome Kinetics in Development, Cell. Rep., 14, 632-647.

65. Haramoto, Y., Oshima, T., Takahashi, S., and Ito, Y. 2014, Characterization of the insulin-like growth factor binding protein family in *Xenopus tropicalis*, Int. J. Dev. Biol., 58, 705-711.

66. Jiang, P., Nelson, J. D., Leng, N., Collins, M., Swanson, S., Dewey, C. N. et al. 2017, Analysis of embryonic development in the unsequenced axolotl: Waves of transcriptomic upheaval and stability, Dev. Biol., 426, 143-154.

67. Wu, R., Liu, Q., Meng, S., Zhang, P., and Liang, D. 2015, Hox cluster characterization of Banna caecilian (*Ichthyophis bannanicus*) provides hints for slow evolution of its genome, BMC Genomics, 16, 468.

68. Kondo, M., Kondo, M., Yamamoto, T., Takahashi, S., and Taira, M. 2017, Comprehensive analyses of hox gene expression in *Xenopus laevis* embryos and adult tissues, Dev. Growth Differ., 59, 526-539.

69. Wacker, S. A., Jansen, H. J., McNulty, C. L., Houtzager, E., and Durston, A. J. 2004, Timed interactions between the Hox expressing non-organiser mesoderm and the Spemann organiser generate positional information during vertebrate gastrulation, Dev. Biol., 268, 207-219.

70. Putnam, N. H., Butts, T., Ferrier, D. E., Furlong, R. F., Hellsten, U., Kawashima, T. et al. 2008, ‘The amphioxus genome and the evolution of the chordate karyotype’, Nature, 453, 1064-1071.

71. Smith, J. J., Putta, S., Zhu, W., Pao, G. M., Verma, I. M., Hunter, T. et al. 2009, Genic regions of a large salamander genome contain long introns and novel genes, BMC Genomics, 10, 19.

72. Sun, C., Shepard, D. B., Chong, R. A., López Arriaza, J., Hall, K., Castoe, T. A. et al. 2012, LTR retrotransposons contribute to genomic gigantism in plethodontid salamanders, Genome Biol. Evol., 4, 168-183.

73. Gardiner, D.M., Blumberg, B., Komine, Y., and Bryant, S.V. 1995, Regulation of HoxA expression in developing and regenerating axolotl limbs, Development, 121, 1731-1741.

74. Endo, T., Tamura, K., and Ide, H. 2000, Analysis of gene expressions during Xenopus forelimb regeneration, Dev. Biol., 220, 296-306.

75. Beck, C.W., Christen, B., and Slack, J.M. 2003, Molecular pathways needed for regeneration of spinal cord and muscle in a vertebrate, Dev. Cell., 5, 429-439.

76. Satoh, A., Endo, T., Abe, M., Yakushiji, N., Ohgo, S., Tamura, K. et al. 2006, Characterization of Xenopus digits and regenerated limbs of the froglet, Dev. Dyn., 235, 3316-3326.

77. Lee-Liu, D., Sun, L., Dovichi, N. J., Larraín, J. 2018, Quantitative Proteomics After Spinal Cord Injury (SCI) in a Regenerative and a Nonregenerative Stage in the Frog, Mol. Cell. Proteomics, 17, 592-606.

